# The stronger downregulation of *in vitro* and *in vivo* innate antiviral responses by a very virulent strain of infectious bursal disease virus (IBDV), compared to a classical strain, is mediated, in part, by the VP4 protein

**DOI:** 10.1101/2019.12.17.879437

**Authors:** KL Dulwich, AS Asfor, AG Gray, ES Giotis, MA Skinner, AJ Broadbent

## Abstract

IBDV is economically important to the poultry industry. Very virulent (vv) strains cause higher mortality rates than other strains for reasons that remain poorly understood. In order to provide more information on IBDV disease outcome, groups of chickens (n=18) were inoculated with the vv strain, UK661, or the classical strain, F52/70. Birds infected with UK661 had a lower survival rate (50%) compared to F52/70 (80%). There was no difference in peak viral replication in the bursa of Fabricius (BF), but the expression of chicken IFNβ, MX1 and IL-8 was significantly lower in the BF of birds infected with UK661 compared to F52/70 (p<0.05) as quantified by RTqPCR, and this trend was also observed in DT40 cells infected with UK661 or F52/70 (p<0.05). The induction of expression of type I IFN in DF-1 cells stimulated with polyI:C (measured by an IFN-β luciferase reporter assay) was significantly reduced in cells expressing ectopic VP4 from UK661 (p<0.05), but was higher in cells expressing ectopic VP4 from F52/70. Cells infected with a chimeric recombinant IBDV carrying the UK661-VP4 gene in the background of PBG98, an attenuated vaccine strain that induces high levels of innate responses (PBG98-VP4^UK661^) also showed a reduced level of IFNα and IL-8 compared to cells infected with a chimeric virus carrying the F52/70-VP4 gene (PBG98-VP4^F52/70^), and birds infected with PBG98-VP4^UK661^ also had a reduced expression of IFNα in the BF compared to birds infected with PBG98-VP4^F52/70^. Taken together, these data demonstrate that UK661 induced the expression of lower levels of anti-viral type I IFN and proinflammatory genes than the classical strain *in vitro* and *in vivo* and this was, in part, due to strain-dependent differences in the VP4 protein.

## Introduction

Infectious bursal disease virus (IBDV) is a highly contagious, immunosuppressive virus belonging to the *Birnaviridae* family [1]. The virus is non-enveloped, with a bi-segmented double stranded (ds) RNA genome encoding 3 open reading frames (ORFs) which are translated and processed to produce 5 viral proteins (VP1-5). Ranking among the top five infectious problems of chickens [2], IBDV poses a continuous threat to the poultry industry though economic losses and welfare concerns. Moreover, as the virus has a preferred tropism for B cells, the majority of which reside in the bursa of Fabricius (BF), surviving birds are often immunosuppressed, less responsive to vaccination programmes, and more susceptible to secondary infections [3, 4].

Disease severity depends on numerous factors including the age and breed of the bird, and the virulence of the infecting IBDV strain [5]. Since the first identification of IBDV in the 1960s, classical (c) strains have circulated worldwide, however, in the 1980s, so-called “very virulent” (vv) strains emerged, complicating IBDV control efforts [6, 7]. The vvIBDV strains cause a far higher mortality rate than classical strains, reaching up to 60-70% in some flocks [8]. However, the molecular basis for the difference in disease outcome remains poorly understood, although it has been demonstrated that both segments A and B contribute to virulence [9]. Segment A encodes the non-structural protein, VP5, and a polyprotein (VP2-VP4-VP3) which is co-translationally cleaved by the protease, VP4 [10]. VP2 is the capsid protein and VP3 is a multifunctional scaffolding protein that binds the genome. The single ORF on Segment B encodes VP1, the RNA-dependent RNA polymerase.

The innate immune response to IBDV infection is characterised by the production of type I IFN responses, including the upregulation of IFNα and IFNβ, that lead to the induction of interferon stimulated genes (ISGs), including MX1, which is one of the top ISGs identified in chicken cells ranked by fold change [11]. The IFN-response aims to provide an antiviral state in infected and bystander cells. In addition, pro-inflammatory cytokines, for example IL-6, IL-8 and IL-1β are produced following IBDV infection that recruit immune cells into the infected bursa of Fabricius (BF) [12–15].

We sought to identify IBDV virulence determinants in order to better understand the molecular basis of phenotypic differences between vv- and c-IBDV strains. Here we report that vvIBDV UK661 down-regulated the expression of antiviral type I IFN responses and pro-inflammatory cytokines compared to cIBDV F52/70 *in vitro* and *in vivo*. Moreover, we demonstrate that the differences in IFN antagonism were, in part, due to strain-dependent differences in the VP4 proteins.

## Materials and methods

### Cells and viruses

DF-1 cells (chicken embryonic fibroblast cells, ATCC number CRL-12203) were sustained in Dulbecco’s modified Eagle’s medium (DMEM) (Sigma-Aldrich, Merck, UK) supplemented with 10% heat inactivated foetal bovine serum (hiFBS) (Gibco, Thermo Fisher Scientific, UK) (Complete DMEM). DT40 cells (immortalised chicken B cell line [16]) were maintained in Roswell Park Memorial Institute (RPMI) media supplemented with L-glutamine (Sigma-Aldrich, Merck), sodium bicarbonate (Sigma-Aldrich, Merck) 10% hiFBS, tryptose phosphate broth (Sigma-Aldrich, Merck) sodium pyruvate (Gibco) and 50mM beta-mercaptoethanol (B-ME) (Gibco) (complete RPMI media). A vv-strain, UK661 [7], and a c-strain, F52/70 [17] of IBDV were kind gifts from Dr Nicolas Eterradossi, ANSES, France. Stocks of both viruses were generated by inoculating 3 week old specific pathogen free (SPF) Rhode Island Red (RIR) chickens and harvesting the BF at 3 days post-infection. BF tissue from 18 birds was pooled, homogenised, and the homogenate mixed with Vertrel XF (Sigma-Aldrich, Merck) and centrifuged at 1,200g for 30 mins. The resulting aqueous phase was harvested and frozen at −80°C. The cell-culture adapted vaccine strain, PBG98, and chimeric viruses within the PBG98 backbone (PBG98-VP4^UK661^ and PBG98-VP4^F52/70^) were propagated in DF-1 cells. Briefly, flasks of DF-1 cells were inoculated with viruses and incubated until cytopathic effect (cpe) was observed, whereupon the supernatant was harvested, centrifuged at 1,200rpm to pellet debris, and the supernatant was aliquoted and frozen at −80C.

### Virus titration by TCID_50._

In order to titrate the UK661 and F52/70 viruses, 96-well U-bottomed plates (Thermo Fisher Scientific) were seeded with the immortal B cell line, DT40, at a seeding density of 1×10^4^ cells per well in 180µL media. A ten-fold dilution series of the UK661 or F52/70 viruses was added to the cells, with 20µL of each dilution added to each well in quadruplicate. Cells were incubated at 37°C for five days, fixed in 4% paraformaldehyde and stained with a primary mouse monoclonal antibody raised against IBDV VP2 [18] and a secondary goat anti-mouse antibody conjugated to Alexa Fluor 488 (Thermo Fisher Scientific). Wells were marked positive or negative for the presence or absence of virus by immunofluorescence microscopy and the TCID_50_/mL calculated by the Reed and Muench method [19]. In order to titrate the PBG98 and chimeric viruses, 96-well plates were seeded with DF-1 cells at a density of 1×10^4^ cells per well in 180µL media. A ten-fold dilution series of the PBG98 or chimeric viruses were added to the cells, with 20µL of each dilution added to each well in quadruplicate. Cells were incubated at 37°C for five days, and wells were marked positive or negative for the presence or absence of cpe and the TCID_50_/mL calculated by the Reed and Muench method [19].

### F52/70 and UK661 in vivo study

Forty-two SPF RIR chickens of mixed gender were obtained from the National Avian Research Facility (NARF) and reared at The Pirbright Institute. Chickens were randomly designated into mock-infected (n=6), F52/70-infected (n=18) and UK661-infected (n=18) groups. At three weeks of age, birds were inoculated with either PBS or a virus dose of 1.8×10^3^ TCID_50_/bird, delivered intranasally, 50 µl per nares. Clinical scores were recorded at least twice daily according to a points-based scoring system (Supplementary Figure 1) that characterised disease as mild (1-7), moderate (8-11), or severe (12-17). The scoring system was developed at The Pirbright Institute and approved by the Home Office (Project Licence number 7008981). Briefly, birds were scored on their appearance, behaviour with and without provocation and handling, and included an assessment of the wattles, combs, feathers, eyes, posture, breathing, interactions with the rest of the flock, ability to evade capture, weight, and crop palpation. Six birds from each infected group were humanely culled at the time points indicated, or when the humane end point (a score of 11) was reached, and tissues were harvested at post-mortem for downstream analysis. The BF was harvested from each bird and divided into two sections, one stored in RNAlater (Thermo Fisher Scientific) for RNA extraction and one snap frozen on dry ice for virus titration by TCID_50_. All animal procedures conformed to the United Kingdom Animal (Scientific Procedures) Act (ASPA) 1986 under Home Office Establishment, Personal and Project licences, following approval of the internal Animal Welfare and Ethic Review Board (AWERB) at The Pirbright Institute.

### PBG98, PBG98-VP4^F52/70^ and PBG98-VP4^UK661^ *in vivo* study

Seventy-two SPF RIR chickens of mixed gender were obtained from the NARF and reared at The Pirbright Institute. Chickens were randomly divided into mock-infected (n=18), recombinant wild-type (wt) PBG98-infected (n=18), PBG98-VP4^F52/70^-infected (n=18) and PBG98-VP4^UK661^-infected (n=18) groups. At three weeks of age, birds were inoculated with either PBS or a virus dose of 1.8×10^3^ TCID_50_/bird, delivered intranasally, 50 µl per nares. Clinical scores were recorded at least twice daily according to the points-based scoring system (Supplementary Figure 1). Six birds from each infected group were humanely culled at 2, 4 and 14 days post-infection and the BF was harvested from each bird and divided into two sections, one stored in RNAlater (Thermo Fisher Scientific) for RNA extraction and one snap frozen on dry ice. All animal procedures conformed to the United Kingdom ASPA 1986 under Home Office Establishment, Personal and Project licences, following approval of the internal AWERB at The Pirbright Institute.

### RNA extraction, RT and qPCR

RNA was extracted from 30mg of homogenised bursal tissue, or from cells in tissue culture wells, using the Monarch Total RNA Miniprep Kit (NEB) according to manufacturer’s instructions. Complementary DNA (cDNA) was generated using SuperScript III Reverse Transcriptase (Invitrogen). The reaction constituents and conditions were consistent with the manufacturer’s instructions. For virus quantification, qPCR was performed using TaqMan™ Universal qPCR Master Mix (Applied Biosystems) according to manufacturer’s instructions. Amplification and detection of targeted genes was performed with the QuantStudio™ 5 (Applied Biosystems) with the following cycling conditions: 50°C for 5 min, 95°C for 2 min, 40°C cycles of 95°C for 3 sec and 60°C for 30 sec. For quantification of host genes, a SYBR green qPCR was performed using Luna® Universal qPCR mix (NEB) according to manufacturer’s instructions. Amplification and detection of targeted genes was performed with the QuantStudio™ 5 (Applied Biosystems) with the following cycling conditions: 95°C for 20 sec, 40 cycles of 95°C for 1 sec and 60°C for 20 sec, then a melt curve step at 95°C for 1 sec, 60°C for 20 sec and 95°C for 1 sec. Primers for all qPCR reactions can be found in Supplementary Table 1. CT values were first normalised to a housekeeping gene (RPLPO) and then to mock inoculated controls and expressed as fold change in a ΔΔCT analysis.

### *In vitro* IBDV infection

DF-1 cells were seeded into 24 well plates at 1.5×10^5^/well and incubated at 37°C overnight to allow adhesion. Virus stocks were diluted in complete DMEM media to the specified multiplicity of infection (MOI) and added to the cells. DT40 cell suspensions were counted, pelleted and resuspended in a solution of virus diluted in complete RPMI media at the specified MOI. Cells were incubated for 1 hour at 37°C and 5% CO_2_. After incubation, the inoculum was removed from the cells and cells were washed with fresh media before incubation in fresh media at 37°C and 5% CO_2_ until the desired time point.

### Luciferase assays for IFNβ

VP4 expression plasmids were designed and synthesised with the gene encoding a chicken codon-optimised enhanced (e) GFP tag at the 5’ end of the VP4 nucleotide sequence. DF-1 cells were seeded into a 24 well plate at a density of 1.5×10^5^ cells/ well and incubated for 24 hours at 37°C and 5% CO_2_ until 80% confluency. Cells were transfected with expression plasmids using Lipofectamine™ 2000 (Invitrogen) according to the manufacturer’s instructions. A final concentration of 500ng plasmid per well was used following optimisation experiments. Briefly, cells were co-transfected with 40ng Renilla luciferase, 80ng of a pGL3 Luciferase reporter plasmid containing the promoter regions of IFNβ upstream of a Firefly luciferase gene to measure type I IFN induction (a kind gift from Steve Goodbourn of St George’s, University of London) and 500ng of eGFP or eGFP-tagged VP4 expression plasmids. Twenty four hours post-transfection, transfection efficiency was confirmed by immunofluorescence microscopy, and protein was extracted from parallel wells for western blot analysis. IFNβ production was stimulated by transfecting cells with (10 μg/mL) polyinosinic: polycytidylic acid (poly I:C) (Sigma Aldrich, Merck). Cells were lysed 6 hours after poly I:C transfection using 100μL 1X passive lysis buffer (Promega). Plates were frozen for at least 30 minutes at −80°C before thawing and reading on a GloMax Multi plate reader (Promega). Briefly, once thawed, 10µL of the sample was added in triplicate to a 96 well opaque white plate (Pierce) and analysed on the plate reader using Stop and Glo reagents (Promega) according to the manufacturer’s instructions. Firefly and Renilla values were recorded and Firefly luciferase values were normalised to Renilla values.

### Western Blot

Cells were lysed in Laemmli Sample buffer (BIO-RAD) containing B-ME, heated to 95°C for 5 minutes, and disrupted by sonication. Samples were subject to sodium dodecyl sulfate-polyacrylamide gel electrophoresis (SDS-PAGE) using a Mini-PROTEAN tetra vertical electrophoresis chamber (BIO-RAD) and transferred to a nitrocellulose membrane using a Trans-Blot Turbo Transfer System (BIO-RAD). Membranes were stained with rabbit anti-VP4 (a gift from Jóse Castón of Centro Nacional de Biotecnología (CNB-CSIC)) and mouse anti-βactin (Thermo Fisher Scientific), followed by donkey anti-rabbit-680 and donkey anti-mouse-800 (LI-COR) and imaged with an Odyssey CLx (LI-COR).

### Generation of chimeric VP4 viruses

A reverse genetics system for the cell-adapted (ca-)IBDV vaccine strain PBG98 was developed in house. Briefly, segment A and segment B were both flanked by the hepatitis delta and hammerhead ribozymes, and cloned into a pSF-CAG-KAN plasmid (Sigma-Aldrich, Merck). These were co-transfected into DF-1 cells with Lipofectamine™ 2000 and the supernatant containing virus was passaged onto additional DF-1 cells to generate viral stocks. Genestrings were synthesised for the PBG98 segment A, replacing the VP4 sequence with that of UK661 or F52/70 (PBG98-VP4^UK661^ and PBG98-VP4^F52/70^, respectively) (GeneArt, Thermo Fisher). Genestrings were digested using KpnI and NheI and ligated into the pSF-CAG-KAN vector. The segment A plasmids PBG98-VP4^UK661^ or PBG98-VP4^F52/70^ were co-transfected into DF-1 cells with the plasmid encoding PBG98 Segment B, and the cultures incubated until cpe was observed. The virus-containing supernatants were passaged onto additional DF-1 cells to generate viral stocks. The sequence of the VP4 gene was determined by Sanger sequencing for all recovered recombinant viruses.

### Statistical Analysis

Statistical significance between experimental groups was determined following a Shapiro-Wilk normality test to confirm whether the data followed a normal distribution for parametric or non-parametric testing. A combination of a two-tailed unpaired Student’s t-test, a Kruskal-Wallis test with a Dunn’s multiple comparison test, or a one-way ANOVA with Tukey’s multiple comparison test on fold change values were performed, using Minitab (v.18).

## Results

### UK661 was more virulent than F52/70 but both replicated to the same *in vivo* peak titre

Groups of 18 chickens were either inoculated with UK661 or F52/70, and 6 chickens were mock-inoculated with PBS alone. Birds were assessed clinically at least twice daily, and humanely culled when humane end-points were reached. At 24 and 48 hours post-infection (hpi), 6 birds per infected group were humanely culled and the BF harvested for quantification of viral replication and host gene expression. At 54 hpi, 3 of the remaining 6 (50%) birds inoculated with UK661 reached their humane end-points and were humanely culled, compared to 1/6 (17%) of birds inoculated with F52/70 (Figure 1A), consistent with UK661 being more virulent than F52/70, as expected. The remaining infected and mock-inoculated birds were humanely culled at 72 hpi. There was no statistically significant difference in the BF: body weight ratio (BF:BW) between groups of birds (Supplementary Figure 2), a metric that is sometimes used as a surrogate of bursal pathology [20, 21]. Moreover, the kinetics of disease progression was similar between the two viral strains, peaking at 54 hpi (Figure 1B), and there was no significant difference in the fold change in viral transcripts measured by RTqPCR between the two strains at any of the time points measured (Figure 1C). We also determined the viral titres in the bursal tissue at each time point by TCID_50_, and although we found UK661 replicated to a lower titre than F52/70 at 24 hpi (*P<0.05), there was no significant difference in viral replication at later time points, which peaked at approximately 8 log_10_ TCID_50_/g of bursal tissue for both strains (Figure 1D).

**Figure 1:**
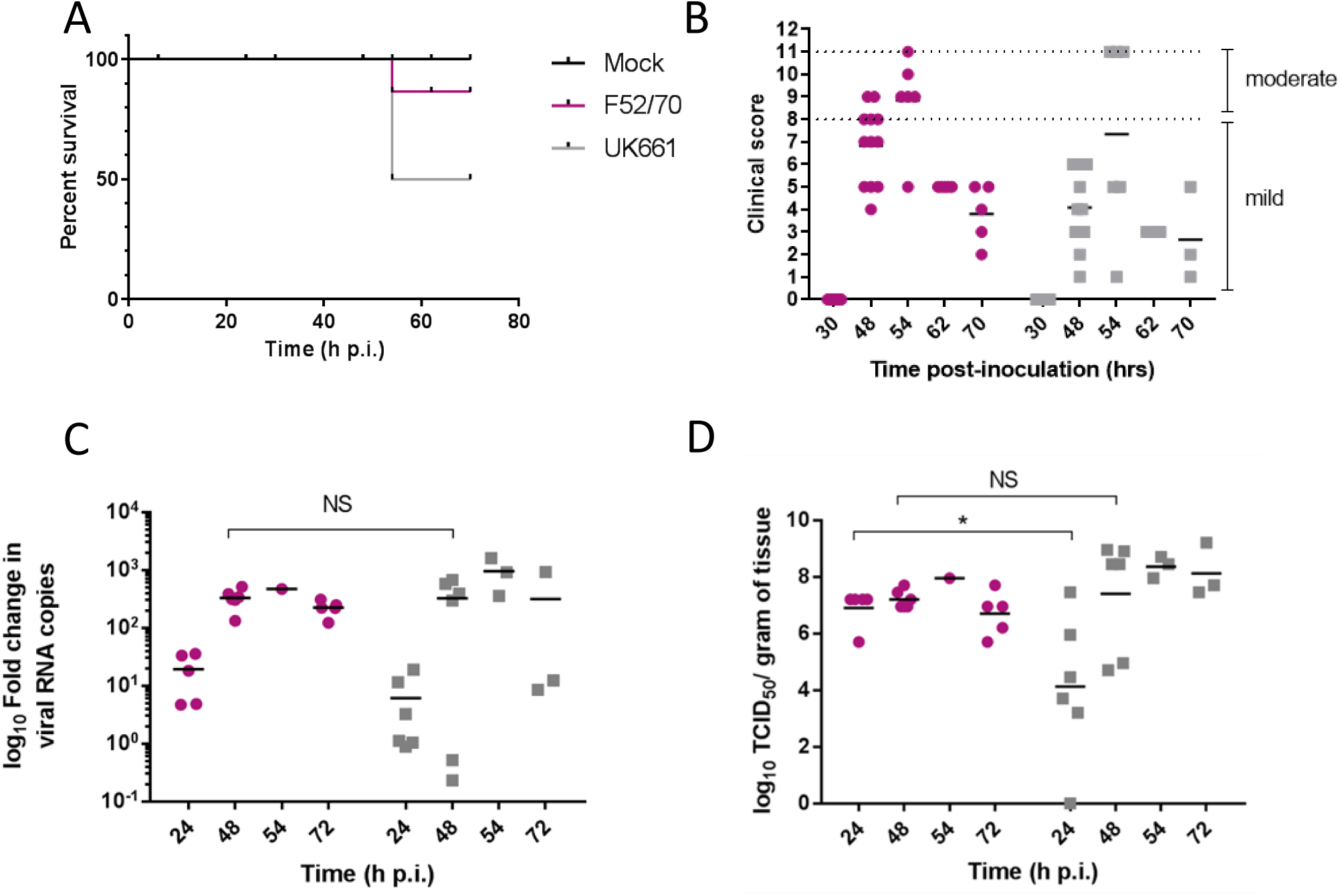
The UK661 strain was more virulent than the F52/70 strain, but both strains replicated to the same peak titre *in vivo*. Birds were checked twice daily by two independent observers for clinical signs and a Kaplan Meier survival curve plotted of mock- (black), F52/70- (pink) and UK661- (grey) inoculated birds that reached their humane end points (clinical score of 11) (A). Clinical signs were quantified by a scoring system and divided into mild (1-7) and moderate (8-11). Each bird was assigned a clinical score at the indicated time points post-infection (B). Six birds per group were humanely culled at 24 and 48 hours post-infection (hpi), one F52/70 and three UK661-infected birds reached their humane end-points at 54 hpi and the remaining birds were culled at 72 hpi. The bursa of Fabricius was harvested at necropsy and the log_10_ fold change in viral RNA copies/g tissue determined by RT-qPCR (C). The infectious titre was determined by titration onto DT40 cells in the method described by Reed & Muench. Virus titres were expressed as log_10_ TCID_50_/g of tissue (D). The horizontal lines are the mean values. Data passed a Shapiro-Wilk normality test before analysis using a two-tailed unpaired Student’s t-test (*P<0.05).

### The expression of type I IFN and pro-inflammatory genes was significantly reduced in BF tissue harvested from birds infected with strain UK661 compared to strain F52/70 *in vivo*

RNA was extracted from BF samples and reverse transcribed to cDNA that was used as the template in qPCR assays targeting chicken type I IFN genes IFNα, IFNβ and Mx1, and pro-inflammatory cytokines IL-6, IL-8 and IL-1β. These genes were selected as they are the most relevant to studying antiviral type I IFN and pro-inflammatory responses, and have previously been shown to be upregulated following IBDV infection [11–15]. The mean expression of IFNα (Figure 2A), IFNβ (Figure 2B), and Mx1 (Figure 2C) was lower in BF samples harvested from UK661-infected birds compared to F52/70-infected birds at 24 and 48 hours post-inoculation, which was statistically significant for IFNβ at 48 hpi and Mx1 at 24, 48, and 72 hpi (*P<0.05). There was no significant difference in the mean expression of IL-1β (Figure 2D) or IL-6 (Figure 2E) between birds infected with the two strains, but IL-8 was significantly reduced in BF samples harvested from UK661-infected birds compared to F52/70-infected birds at 48 and 72 hours post-inoculation (*P<0.05) (Figure 2F). Taken together, these data demonstrate that the expression of antiviral type I IFN responses (IFNβ and MX1) and pro-inflammatory cytokine responses (IL-8) in the BF was reduced following infection with the vv strain compared to the classical strain.

**Figure 2:**
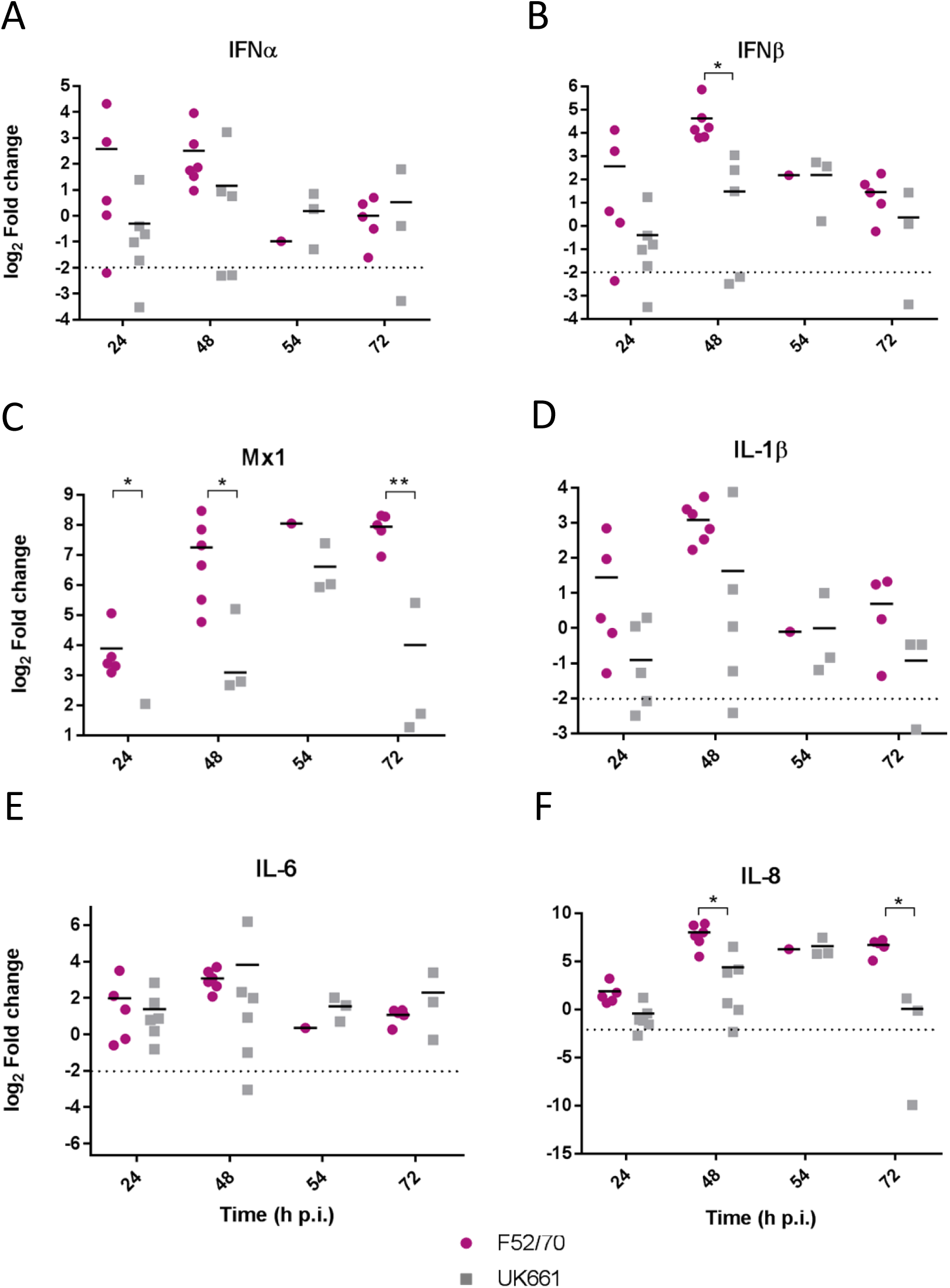
The expression of type I IFN and pro-inflammatory genes was significantly reduced in BF tissue harvested from birds infected with strain UK661 compared to strain F52/70 *in vivo*. The bursa of Fabricius was harvested from mock and infected birds at necropsy and RNA extracted. RNA was reverse transcribed and amplified by quantitative PCR using specific primer sets for target genes. The CT values were normalised to the housekeeping gene RPLPO and the log_2_ fold change in gene expression determined for the infected samples relative to the mock-infected samples in a ΔΔCT analysis and plotted for individual birds. The horizontal lines are the mean values. Data are representative of at least three replicate experiments and passed a Shapiro-Wilk normality test before analysis using a two-tailed unpaired Student’s t-test (*P<0.05, **P<0.01). The dashed horizontal line represents the cut-off, below which genes were significantly down-regulated compared to mock-inoculated birds.

### The expression of type I IFN and pro-inflammatory genes was significantly reduced in B cells infected with UK661 compared to F52/70 *in vitro*

We compared the replication of UK661 and F52/70 and the expression of type I IFN and proinflammatory cytokines in DT40 cells, an immortalised avian B cell-line. Cells were infected with UK661 or F52/70 (MOI 0.1) and the expression of virus and host-cell transcripts quantified by RTqPCR at 14, 48 and 72 hpi. UK661 replicated to a significantly higher titre than F52/70 across all time points (***P<0.001) (Figure 3A). However, cells infected with UK661 showed significantly reduced expression of IFNα, IFNβ and Mx1 compared to cells infected with F52/70 at 14, 48 and 72 hpi (*P<0.05, **P<0.01, ***P<0.001) (Figure 3B-D). Expression of IL-1β and IL-6 was also significantly reduced in cells infected with UK661 compared to F52/70 at multiple time-points (*P<0.05, **P<0.01, ***P<0.001) (Figure 3E and F). In contrast, IL-8 expression was significantly higher in cells infected with UK661 than F52/70 at 48 hpi (***P<0.01) (Figure 3G). Taken together, these data suggest that the vv strain is able to reduce the expression of mRNAs for IFNα, IFNβ, Mx1 and the pro-inflammatory cytokines IL-1β and IL-6 to a greater extent than the classical strain *in vitro,* confirming our *in vivo* data.

**Figure 3:**
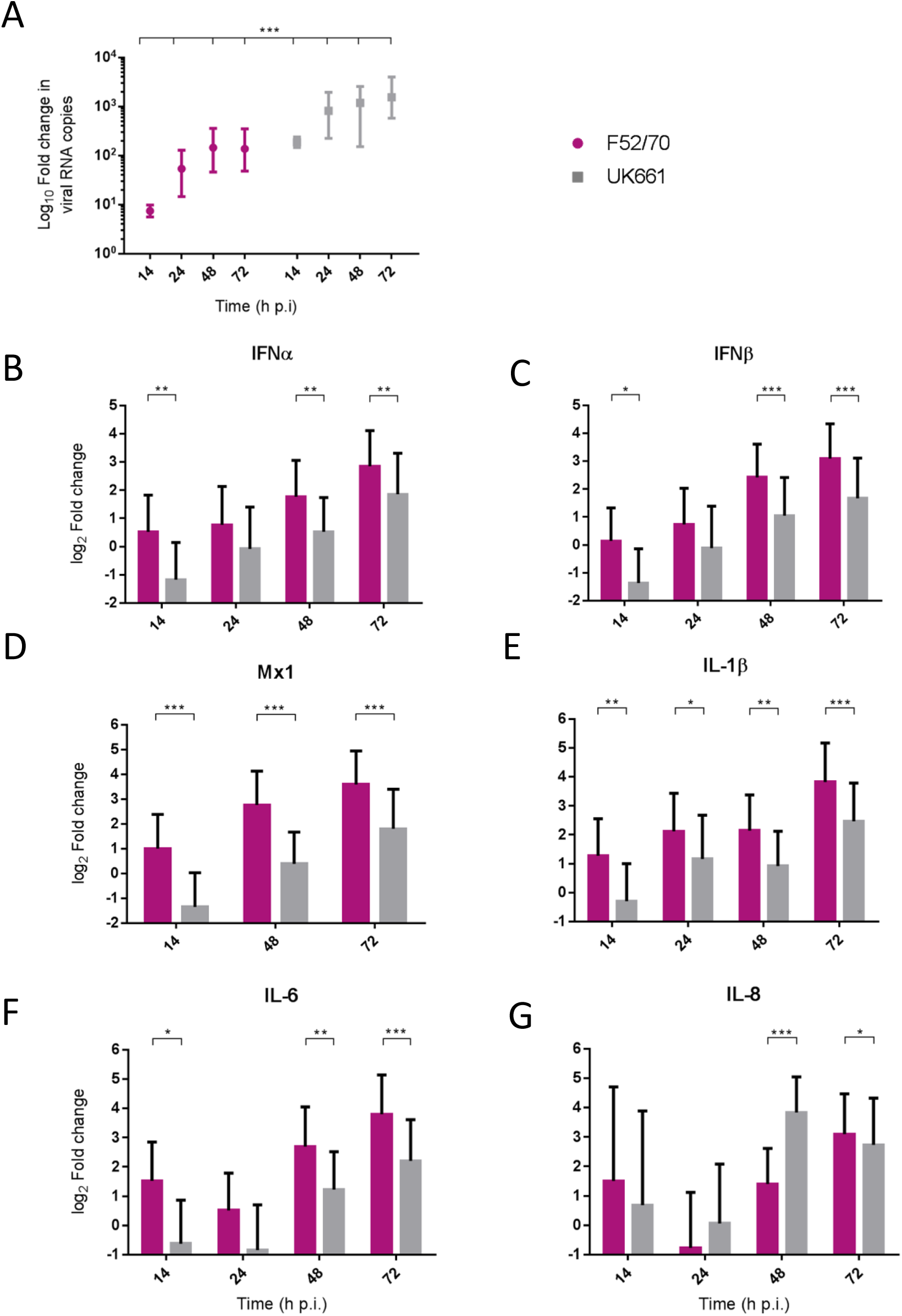
The expression of type I IFN and pro-inflammatory genes was significantly reduced in B cells infected with strain UK661 compared to strain F52/70 *in vitro*. DT40 Cells were infected at an MOI of 0.1 with either the UK661 or F52/70 IBDV strains, or mock-infected with media alone and RNA was extracted from the cells at the indicated time points post-infection. RNA was reverse transcribed and amplified by qPCR using specific primer sets. The CT values were normalised to the housekeeping gene RPLPO and the log_10_ fold change in virus gene expression determined for the infected samples relative to the mock-infected samples in a ΔΔCT analysis and plotted (A). The log_2_ fold change in host-cell gene expression was also determined for the infected samples relative to the mock-infected samples in a ΔΔCT analysis and plotted (B-G). Data subsequently passed a Shapiro-Wilk normality test before being analysed by a one-way ANOVA and a Tukey’s multiple comparison test (*P<0.05) (A), or a two-tailed unpaired Student’s t-test (*P<0.05, **P<0.01, ***P<0.001). Data shown are representative of at least three replicate experiments, columns represent the mean values, and error bars represent the standard deviation of the mean.

### The UK661-, but not F52/70-, VP4 protein antagonised IFNβ induction *in vitro*

The VP4 protein of IBDV has previously been identified as an IFN antagonist [22]. We hypothesised that strain-dependent differences in the VP4 proteins would be responsible for the observed differences in type I IFN responses between the strains. In order to address this, VP4 sequences across groups of very virulent, classical and attenuated/cell-adapted IBDV strains available on the NCBI database were aligned using Clustal Omega [23] (Supplementary Figure 3A). Compared to F52/70 VP4, there were 9 amino acids different in the UK661 VP4 protein (V31I, D114V, D122G, R132K, N141S, C170Y, K175N, P205S, and H241D). Interestingly, 5 of these were also found in other vv strains, from diverse geographical regions, but not in classical or vaccine strains (31I, 170Y, 175N, 205S, and 241D). The VP4 protein structures were modelled (using PyMOL version 2.0, Schrödinger, LLC) to the known VP4 structure of another member of the *Birnaviridae* family, the Yellowtail Ascites Virus (YAV; 4izk.2.A) (Supplementary Figure 3B), as the structure of IBDV VP4 protein remains unsolved at the time of writing. The YAV VP4 sequence shares 24.75% amino acid identity and lacks the last 25 amino acids of the IBDV sequence so the amino acid at position 241 is absent. Nevertheless, based on this model, the amino acid differences between the two VP4 proteins were found to cause some alterations in the predicted secondary structure, for example the presence of β-strands in the UK661 VP4 molecule that are not seen in the F52/70 VP4 molecule (supplementary Figure 3B, dashed boxed regions).

To determine whether the VP4 proteins from UK661 and F52/70 IBDV inhibited IFNβ induction, we co-transfected DF-1 cells with an IFNβ luciferase reporter plasmid and plasmids encoding VP4 proteins from either UK661 or F52/70, tagged with enhanced GFP (eGFP) at the N-terminus (eGFP-UK661-VP4 and eGFP-F52/70-VP4). The eGFP tag was used to monitor transfection efficiency. Transfected cells were stimulated with poly I:C to induce the production of IFNβ that was quantified by measuring Firefly luciferase units normalised to Renilla luciferase. A low level of IFNβ induction was observed in all groups in the absence of poly I:C stimulation, with little difference between those transfected with the VP4 expression plasmids and a plasmid expressing eGFP alone (vector control plasmid) (Figure 4). In contrast, upon stimulation with poly I:C, there was an increase in IFNβ induction in cells expressing eGFP alone that was significantly reduced in cells expressing eGFP-UK661-VP4 (**P<0.01). This reduction was not seen in cells expressing eGFP-F52/70-VP4, despite more protein detected by western blot than eGFP-UK661-VP4. In fact, there was a significant increase in IFNβ induction compared to cells expressing either eGFP-UK661-VP4 or eGFP alone (**P<0.01). These data demonstrate that the VP4 protein from the vv IBDV strain down-regulates IFNβ induction *in vitro*, but the VP4 protein from the classical strain does not.

**Figure 4:**
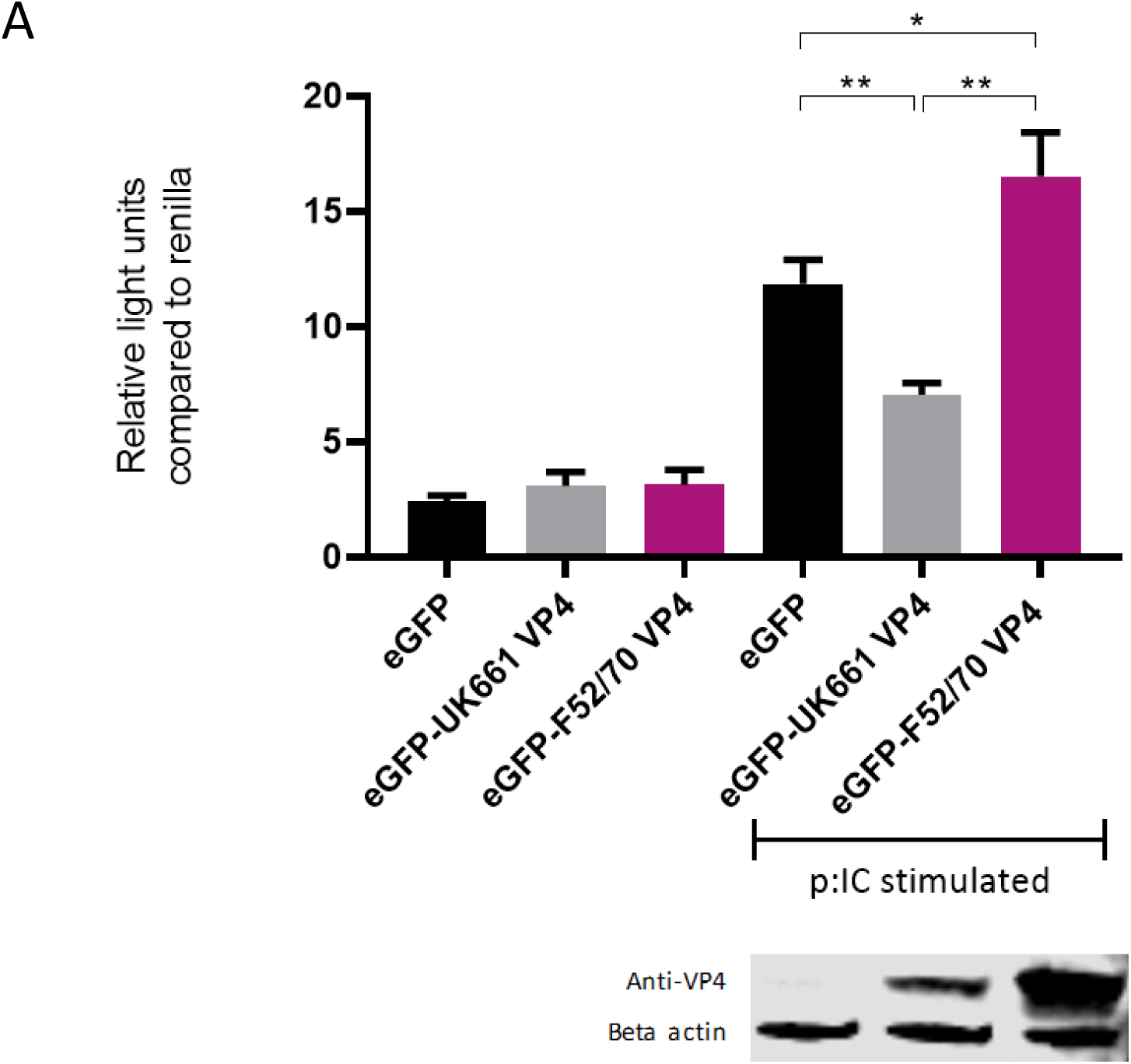
The VP4 protein from the UK661 strain antagonised IFNβ induction, but the VP4 protein from the F52/70 strain did not *in vitro*. DF-1s were transfected with the chicken IFNβ promoter Firefly luciferase reporter and a constitutively active Renilla expression plasmid and 500ng of either eGFP-UK661-VP4 or eGFP-F52/70-VP4 expression plasmids, or a control plasmid expressing eGFP alone. Twenty-four hours post-transfection, cells were re-transfected with poly I:C. At 6 hours post-transfection, cells were lysed and luciferase activity quantified. Firefly luciferase activity was normalised to Renilla expression. Data presented are the means of three independent experiments and passed a Shapiro-Wilk normality test before analysis using a two-tailed unpaired Student’s t-test (*P<0.05, ** P<0.01). Error bars represent the standard error of the mean (SEM). In a parallel experiment, transfected cells were lysed and samples denatured and subject to SDS-PAGE gel electrophoresis followed by transfer to a nitrocellulose membrane and staining with anti-VP4 and anti-β-actin antibodies in a western blot.

### The ability of the UK661 VP4 protein to antagonise type I IFN responses was reduced in the context of the whole virus *in vitro* and *in vivo*

In order to determine the extent to which strain dependent differences in the VP4 protein antagonised type I IFN responses in the context of a whole virus, chimeric viruses were generated with the VP4 gene from either the UK661 or F52/70 strains in the backbone of a highly attenuated cell culture-adapted vaccine strain, PBG98 (PBG98-VP4^UK661^ and PBG98-VP4^F52/70^ respectively). DF-1 cells were infected with recombinant chimeric or wt PBG98 viruses (MOI 1) and RNA was extracted, reverse transcribed and the fold change in viral RNA quantified by qPCR at several time points post-infection. All viruses replicated to a maximum of 10^4^-10^5^ fold change in viral RNA per mL supernatant, with no significant differences in viral replication kinetics (Figure 5A). Consistent with our previous observations, the expression of IFNα was significantly reduced in cells infected with the PBG98-VP4^UK661^ virus compared to cells infected with the PBG98-VP4^F52/70^ virus at 24 hpi (Figure 5B), and, although not statistically significant, the same trend was observed 24 hpi for IFNβ and Mx1 (Figure 5C and D). Interestingly, by 24 hpi, both chimeric viruses induced elevated IFNβ compared to the recombinant wt PBG98 virus. In addition, by 24 hpi, the PBG98-VP4^UK661^ virus induced a lower level of expression of IL-1β and IL-8 than the PBG98-VP4^F52/70^ virus, which reached statistical significance for IL-8 (****P<0.0001). These data demonstrate that although the replication titres of the two chimeric viruses were similar, the expression of type I IFNα and pro-inflammatory cytokine IL-8 was lower during infection with the PBG98-VP4^UK661^ virus compared to the PBG98-VP4^F52/70^ virus.

**Figure 5:**
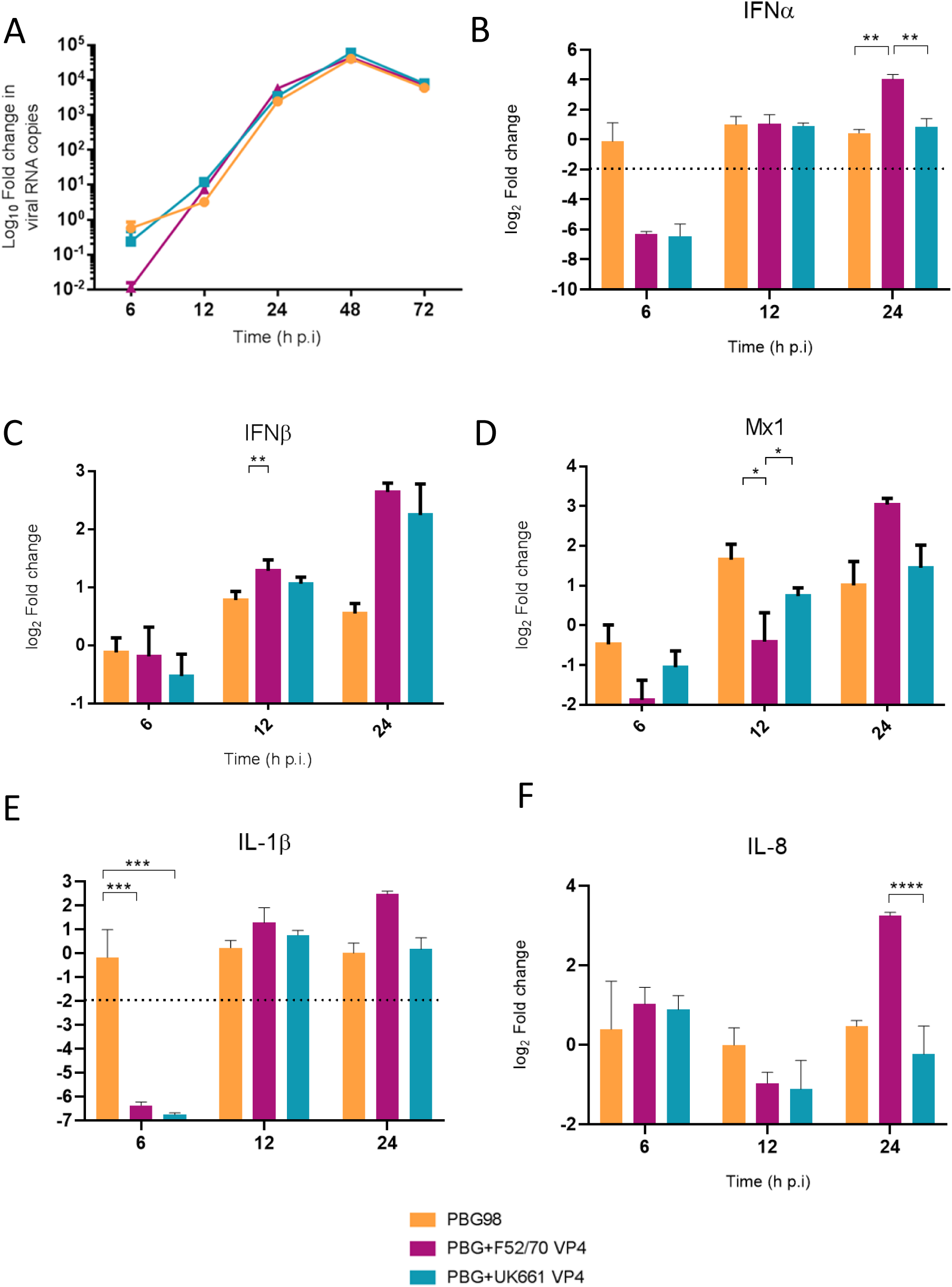
The ability of the UK661 VP4 protein to antagonise type I IFN responses was reduced in the context of the whole virus *in vitro*. DF-1 cells were infected with PBG98, PBG98-VP4^UK661^ and PBG98-VP4^F52/70^ viruses at an MOI of 1, before RNA was extracted at the indicated time points post-infection and reverse transcribed. Virus specific primers were used to amplify the cDNA by quantitative PCR, the CT values were normalised to the housekeeping gene RPLPO and the log_10_ fold change in virus gene expression was determined for the infected samples relative to the mock-infected controls in a ΔΔCT analysis and plotted. A Kruskal-Wallis test was performed with a Dunn’s multiple comparison test where no significant difference was found at any time point between the three viruses (A). A panel of genes, IFNα (B), IFNβ (C), Mx1 (D), IL-1β (E), and IL-8 (F), were amplified by quantitative PCR using specific primer sets for target genes, before the CT values were normalised to the housekeeping gene RPLPO and the log_2_ fold change in gene expression determined for the infected samples relative to the mock-infected controls in a ΔΔCT analysis and plotted. Data are representative of at least three replicate experiments and passed a Shapiro-Wilk normality test before analysis using a two-tailed unpaired Student’s t-test (*P<0.05, **P<0.01, ***P<0.001). The mean values are plotted and the error bars are the standard error of the mean (SEM). The dashed horizontal line represents the cut-off, below which genes were significantly down-regulated.

To compare virus replication kinetics and host gene expression *in vivo,* groups of 18 chickens were inoculated with chimeric and wt PBG98 viruses, or mock inoculated with buffer alone. At 2, 4 and 14 days post-inoculation, the BFs were harvested, RNA was extracted and the expression of IBDV, IFNα, IFNβ, Mx1, IL-6, IL-8, and IL-1β quantified by RTqPCR. There was no significant difference in viral replication between any of the groups (Figure 6A). However, virus replication was somewhat low (up to 10^3^ fold change in viral RNA per gram of BF tissue), possibly due to the cell-culture adapted nature of the backbone. Consistent with our previous observations, at day 2 post-inoculation, the expression of IFNα was significantly lower in the BF of birds infected with the PBG98-VP4^UK661^ virus compared to the PBG98-VP4^F52/70^ virus (*P<0.05) (Figure 6B). This trend was the same at day 4 post-inoculation although this did not reach statistical significance. Likewise, the average IFNβ expression at day 4 post-inoculation was lower in the BF of birds infected with the PBG98-VP4^UK661^ virus compared to the PBG98-VP4^F52/70^ virus, but this did not reach statistical significance (Figure 6C). Mx1 expression was similar in birds infected with PBG98 and the chimeric viruses at days 2 and 4 post-inoculation (Figure 6D). Comparing the pro-inflammatory response between these viruses, IL-1β expression was significantly lower in the BFs of birds inoculated with either of the chimeric viruses at 2 days post-inoculation compared to recombinant wt PBG98 (***P<0.001) (Figure 6E), but there was no significant difference in IL-8 expression between any of the virus groups at day 2 or 4 post-inoculation (Figure 6F). Taken together, these data demonstrate that the chimeric virus containing the VP4 gene from the vvIBDV strain UK661 induced a lower level of type I IFNα compared to the chimeric virus containing the VP4 gene from the cIBDV strain F52/70 both *in vitro* and *in vivo*. However, the effect of the VP4 protein on IFNβ, MX1 or pro-inflammatory cytokines was reduced in the context of virus infection, compared to ectopic expression.

**Figure 6:**
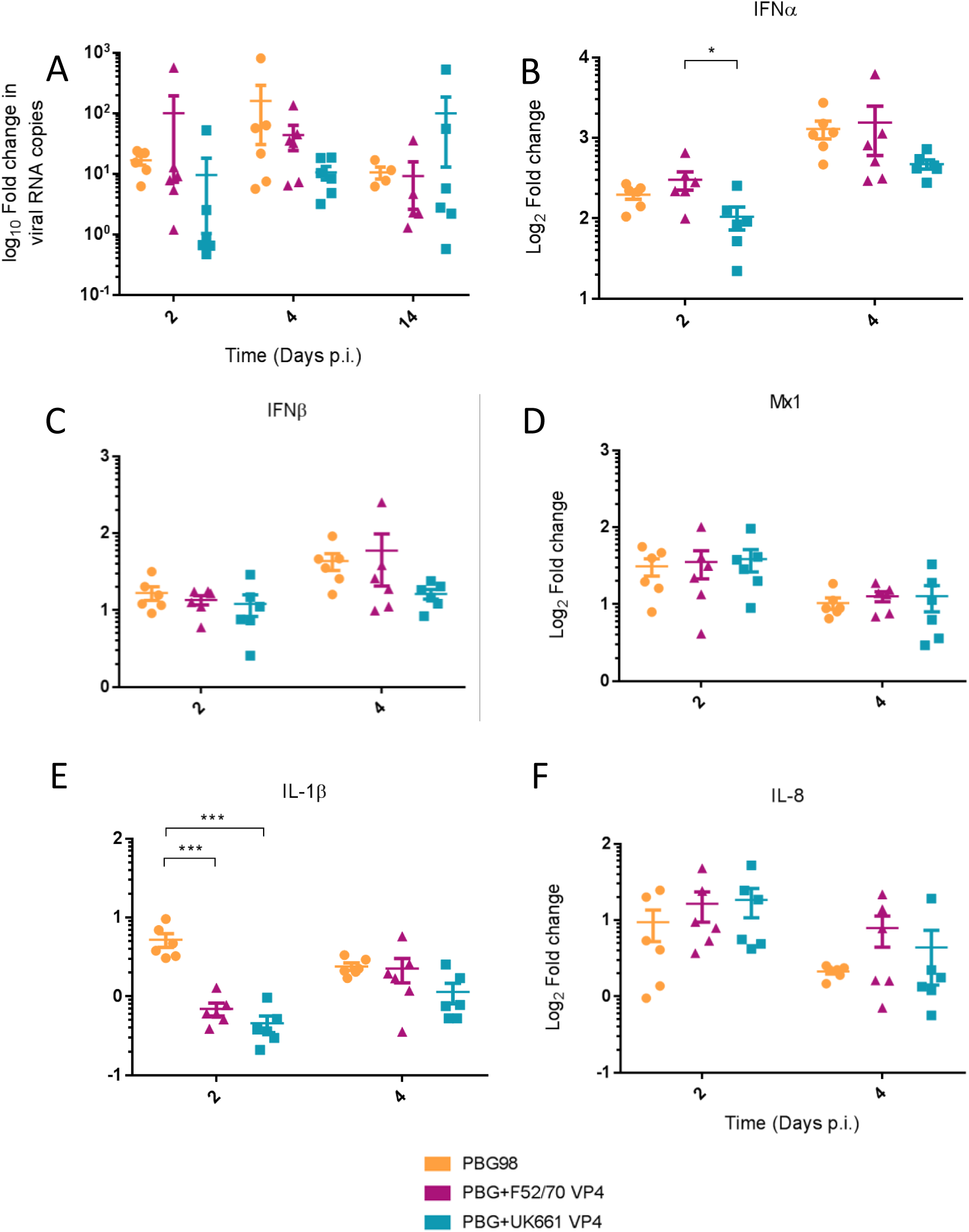
The ability of the UK661 VP4 protein to antagonise type I IFN responses was reduced in the context of the whole virus *in vitro*. Birds were inoculated with 1.8×10^3^ TCID_50_ of the PBG98, PBG98-VP4^UK661^ and PBG98-VP4^F52/70^ viruses, and the bursa of Fabricius was harvested at necropsy from 6 birds per group at 2, 4 and 14 days post-inoculation. RNA was extracted prior to reverse transcription to cDNA and qPCR amplification with virus-specific primers. CT values were normalised to a housekeeping gene and expressed as log_10_fold change viral RNA relative to mock-infected samples as per the ΔΔCT method. The data passed a Shapiro-Wilk normality test before being analysed using a two-way ANOVA (not significant) (A). At 2 and 4 days post-inoculation, cDNA was amplified by qPCR for a panel genes: IFNα (B), IFNβ (C), Mx1 (D), IL-1β (E), and IL-8 (F). The CT values were normalised to the housekeeping gene RPLPO and expressed relative to mock-infected samples using the ΔΔCT method. Data are representative of at least three replicate experiments and passed a Shapiro-Wilk normality test before analysis using a two-tailed unpaired Student’s t-test (*P<0.05, ***P<0.001). Horizontal lines represent the mean and error bars represent the standard error of the mean (SEM).

## Discussion

Very virulent strains of IBDV emerged in the 1980s, causing up to 60% mortality in some commercial flocks [8]. However, the molecular basis for this increased virulence remains poorly understood [24, 25]. We have previously shown that the vvIBDV UK661 was able to down-regulate type I IFN and a selection of ISGs to a greater extent than a vaccine strain, D78, in primary B cells cultured and infected *ex vivo* [26]. Here, we extend these observations by demonstrating that UK661 is also able to down-regulate type I IFN and pro-inflammatory cytokine responses compared to a classical field strain, and we confirm that this occurs not only *in vitro*, but also *in vivo* (Figures 2 and 3).

Other studies have compared strain-dependent differences in innate immune responses to IBDV infection, however, the majority have compared vv strains with cell culture adapted (ca) strains. For example, He *et al.* found TLR3, IL-8 and IFNβ expression were more up-regulated in response to the vvIBDV strain than a vaccine strain, and Liu *et al.* found elevated expression of cytokines following vvIBDV infection compared to a vaccine strain, however the vv strain replicated to significantly higher titres that the vaccine strains in both studies, making comparison of gene expression changes challenging [27, 28]. In contrast, in our study, there was no significant difference in peak virus replication between the F52/70 and UK661 strains, meaning that differences in gene expression are due to something other than the amount of virus present. Unfortunately, some studies comparing the innate immune response following vvIBDV infection to caIBDV strains also inoculated birds with different amounts of virus, making a direct comparison of gene expression difficult [29]. To our knowledge, only one previous study, by Eldaghayes et al., has compared classical and vvIBDV strains *in vivo* [30]. Our data are consistent with this work, which also reported that a vv IBDV strain induced reduced type I IFN responses compared to a classical strain. However, the authors conducted two separate *in vivo* studies, one with each virus, and did not compare the two viruses in the same study. Moreover, birds were inoculated with a different dose of each virus, making a comparison of gene expression challenging. We extend these observations by directly comparing the vv and classical strains in the same *in vivo* study, in birds inoculated with the same dose of each virus.

We also demonstrate that the differences in IFN antagonism are, in part, due to strain-dependent differences in the VP4 proteins (Figures 4, 5 and 6). The VP4 protein from vvIBDV strain Lx has previously been shown to act as an IFN antagonist through an interaction with the host glucocorticoid-induced leucine zipper (GILZ) protein [14, 22]. GILZ plays a key role in the regulation of NF-κB activation by binding to the p65 subunit and preventing its translocation into the nucleus and the downstream expression of cytokines [31]. The Lx VP4 protein has been shown to bind to GILZ, preventing its ubiquitination and degradation, resulting in its accumulation in the cytoplasm. Consequently, this VP4-GILZ interaction enhances the inhibition of p65 translocation into the nucleus, leading to a reduction in the expression of pro-inflammatory cytokines and type I IFN responses [14]. We extend these observations by demonstrating that there are strain dependent differences in the extent to which VP4 antagonises type I IFN induction. Elucidating differences in the mechanism of action of the different VP4 proteins was beyond the scope of this study, however, it might be possible that while the UK661 VP4 is capable of binding GILZ in a manner similar to the Lx VP4, the ability or affinity of F52/70 VP4 to bind GILZ may be reduced. However, additional experiments are needed to confirm this.

We demonstrated the involvement of the VP4 protein in the antagonism of type I IFN responses both by luciferase reporter assay, and the use of chimeric viruses expressing the VP4 protein from either UK661 or F52/70. However, the phenotype we observed with the chimeric viruses was less pronounced than that observed by luciferase reporter assay as, while the PBG98-VP4^UK661^ virus induced lower levels of IFNα, IFNβ and MX1 than the PBG98-VP4^F52/70^ virus *in vitro*, this only reached statistical significance for IFNα at 24 hpi. This suggests that the VP4 gene is not the sole determinant of the difference in type I IFN responses observed between the UK661 and F52/70 viruses. Other IBDV proteins have previously been implicated in the inhibition of type I IFN responses, for example the VP3 protein is known to bind the dsRNA genome of IBDV and competitively inhibit detection by MDA5, preventing the downstream signalling of the IFN pathway and the production of type I IFN (Ye *et al.* 2014), and it is likely that multiple genes work in concert to produce the observed phenotypes.

When the chimeric viruses were inoculated into chickens, both replicated to low titres and caused few clinical signs, most likely because these viruses possess the backbone of a cell culture adapted, highly attenuated virus. We were therefore unable to ascertain whether strain-dependent differences in VP4 affected IBDV virulence. Previous studies, using chimeric viruses with segments A and B from strains of differing virulence, found that both segments contributed to virulence [9, 32–34]. While the mechanism is not yet understood, the effect of VP1 mutations may be related to viral replication, whereas VP2 has been shown to activate apoptosis via the reduction of the anti-apoptotic molecule, ORAOV1 [35], and VP5 plays a key role in apoptosis by preventing it early during infection and by activating it at later time points [36]. Virulence is therefore likely to be a complex phenotype, however, it is possible that the VP4 sequence could contribute to this. Five of the nine amino acid residues in UK661 that were different from F52/70 are also found in diverse vv IBDV strains from different geographical regions, but were not found in other classical or attenuated strains (31I, 170Y, 175N, 205S, and 241D) (Supplementary Figure 3A). It is therefore possible that enhanced ability to antagonise type I IFN responses is feature of vvIBDV strains, and these amino acids represent a VP4 “genetic signature of virulence”. Moreover, given that the classical strains emerged in the 1960s and vv strains subsequently emerged in the 1980s (apparently and probably by segment reassortment), it is tempting to speculate that vvIBDVs evolved to have a VP4 protein with an enhanced ability to inhibit NF-κB activation, promoting increased virus fitness due to suppressed antiviral responses. Consistent with this hypothesis, the replication of the UK661 virus was significantly enhanced in DT40 cells compared to the F52/70 virus (Figure 3A). Interestingly, the clinical scores of birds inoculated with the F52/70 virus were actually higher than birds inoculated with UK661 at 48 hpi (Figure 1 B). At first this seems counter-intuitive that the vv strain would cause less severe symptoms than the classical strain, however, this may be due to the increased expression of type I IFN and pro-inflammatory genes in birds inoculated with the F52/70 virus, as stimulation of these innate immune responses would be expected to result in clinical signs such as lethargy, depression and ruffled feathers as observed to a greater extent in the F52/70-inoculated group. This further underpins the complexity of defining virulence and highlights that our understanding of why some birds reach humane end points, whereas others do not, remains unknown.

Our study is not without limitations: Gene expression was quantified by RTqPCR, and only a small panel of genes were investigated. It would be beneficial in the future to compare gene expression by RNA-Seq to gain a more comprehensive comparison of strain-dependent differences in expression. Never-the less, our dataset does allow us to draw useful conclusions. Additionally, we only compared the UK661 strain with the F52/70 strain and it would be interesting to compare vv, classical, and vaccine strains, and possibly also diverse strains from different geographical regions, or serotype 1 compared to serotype 2. However, this was beyond the scope of the current project. Despite its limitations, our study does provide useful information that can be used to inform IBDV surveillance efforts and improve IBDV vaccines: Identifying genetic signatures of increased IBDV virulence could be used to better inform national surveillance efforts in order to calculate the potential threat of an emerging strain as early as possible. Moreover, identifying genetic signatures of attenuation could be used to engineer a rationally designed live vaccine candidate. For example, it might be beneficial to explore the potential, as novel vaccine candidates, of chimeric viruses engineered with the VP4 gene from attenuated strains in the backbone of a field strain.

Taken together, our data demonstrate that UK661 induced the expression of lower levels of anti-viral type I IFN responses than the classical strain *in vitro* and *in vivo* and this was, in part, due to strain-dependent differences in the VP4 protein. We speculate that this might enhance viral fitness and contribute to the enhanced virulence of UK661. This provides new information that could be used to improve IBDV surveillance efforts and control strategies.

## Acknowledgements

This work was funded by BBS/E/I/00001845, BBS/E/I/00002115, BBS/E/I/00007031, BBS/E/I/00007038, BB/K002465/1 and BBS/E/I/00007039. ESG is also supported by funding from the Wellcome Trust new investigator award (104771/Z/14/Z). We thank Nicolas Eterradossi and Sébastien Soubies of ANSES France for supplying the IBDV strains, Steve Goodbourn of St George’s, University of London for supplying the ch-IFNβ luciferase reporter plasmid, Joe James of APHA for supplying the sequences of the chicken IL-1β, IL-6 and IL-8 primers, and Jóse Castón of Centro Nacional de Biotecnología (CNB-CSIC) for the anti-VP4 antibody. We are also grateful for the husbandry support and assistance from the animal services team at The Pirbright Institute during *in vivo* studies.

## Author Contributions

KD performed experiments, analysed data and wrote the first draft of the manuscript, AA and AG performed experiments, EG and MS helped analyse data and edited drafts of the manuscript, and AB conceptualised the study, obtained funding, helped conduct experiments and analyse data, and edited drafts of the manuscript.

---

**Supplementary Table 1:**
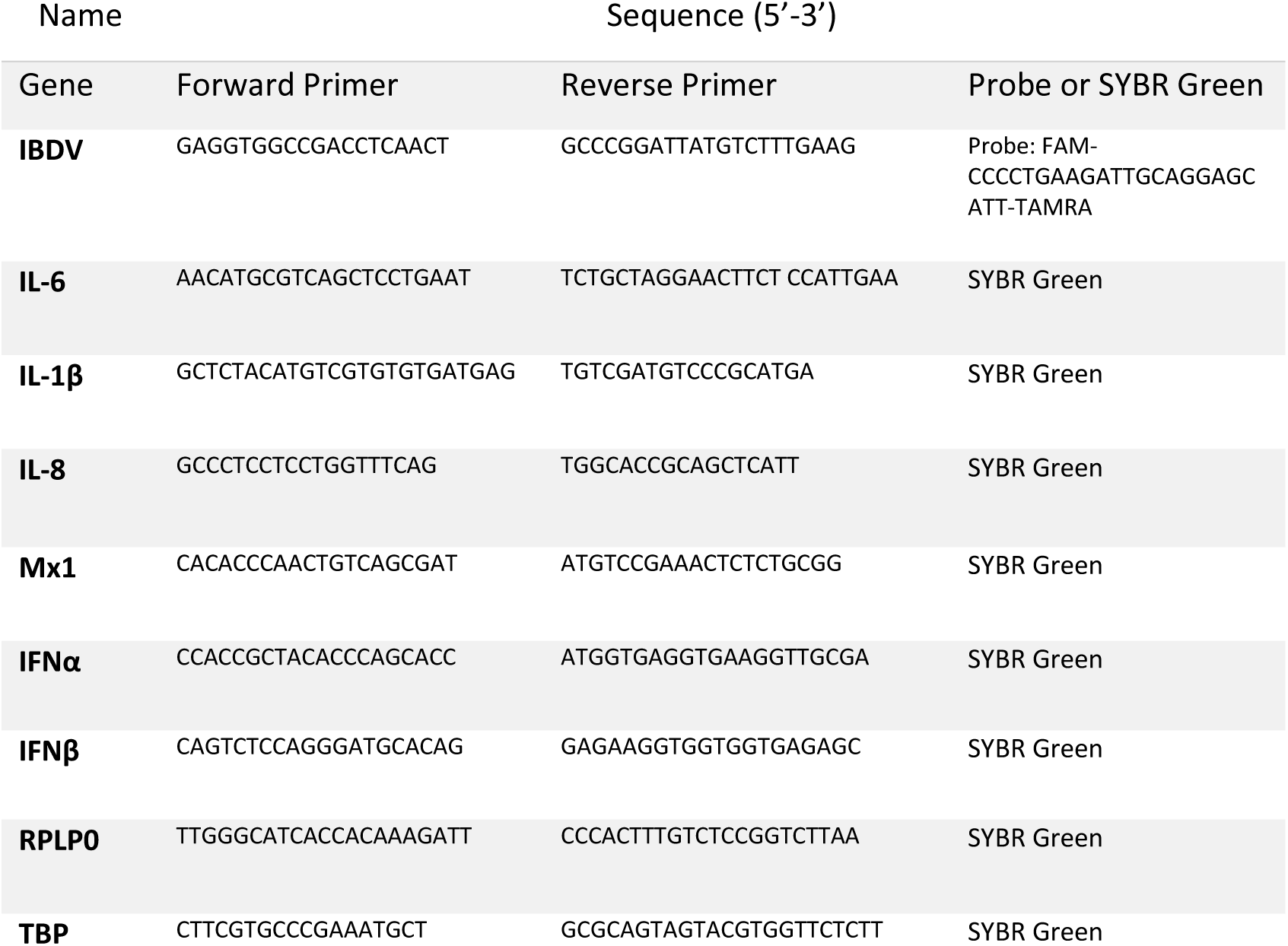
Primers used in this study for qPCR.

**Supplementary Figure 1.**
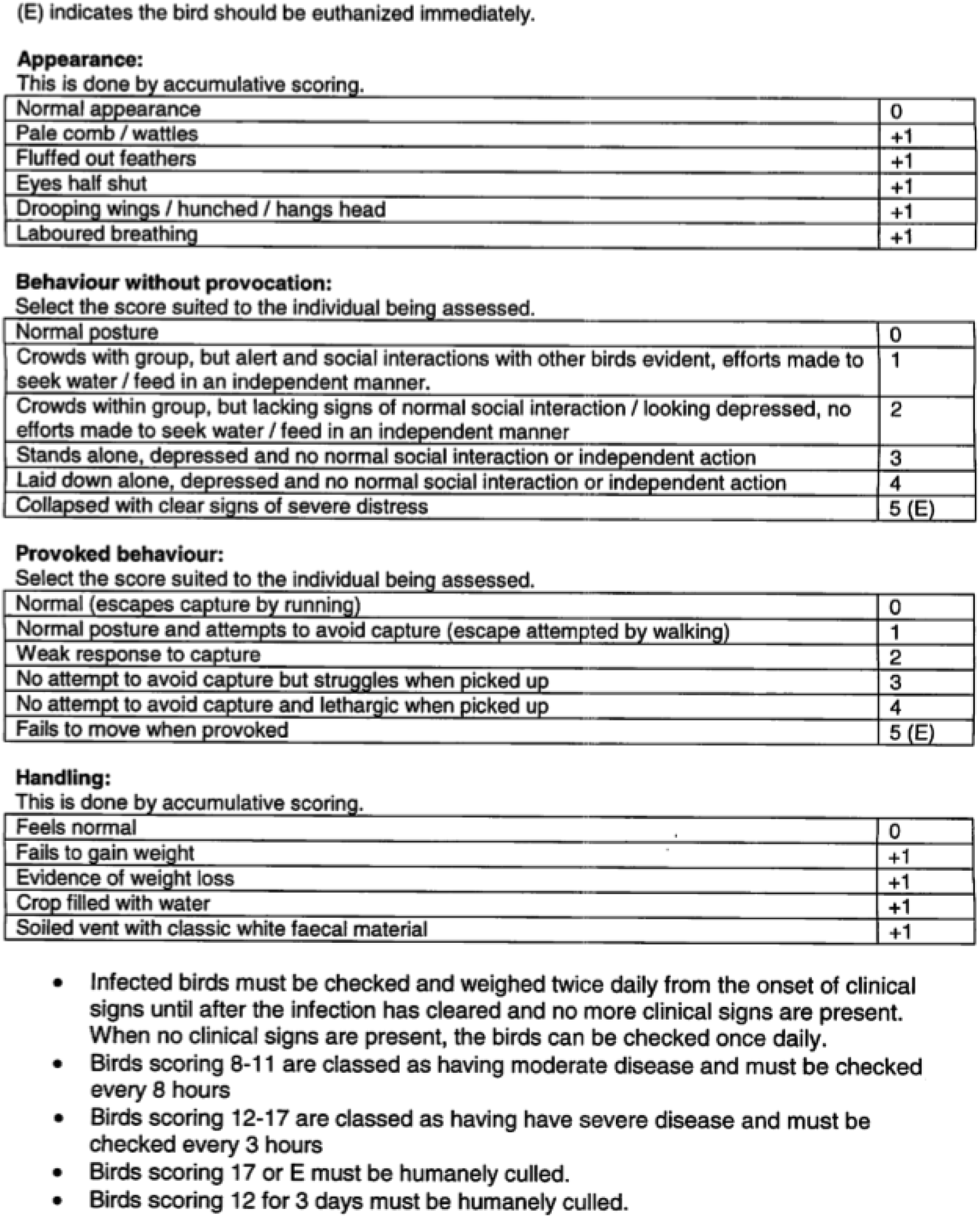
Clinical Scoring system used in *in vivo* studies. Birds were checked at least twice daily following inoculation by two independent observers and the clinical signs scored. Birds were humanely culled when their humane end-points were reached (a clinical score of 11 in this study).

**Supplementary Figure 2:**
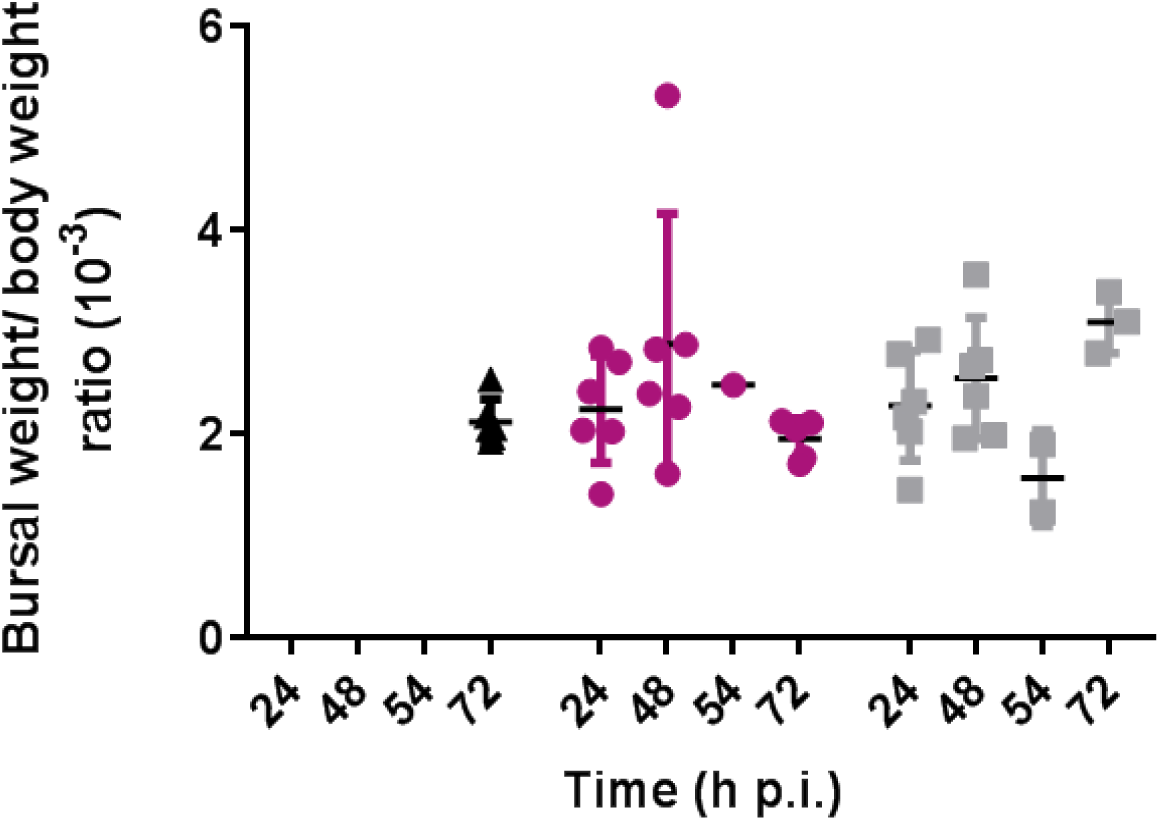
No significant difference was observed between the Bursa of Fabricius: body weight (BF:BW) ratios of mock, F52/70- or UK661-inoculated birds. Six birds per group were humanely culled at 24 and 48 hpi. One bird infected with F52/70 and three birds infected with UK661 were humanely culled at 54 hpi due to reaching their humane end points. The remaining infected birds and mock-inoculated birds were humanely culled at 72 hpi. Birds were weighed and the bursa of Fabricius was removed at necropsy and weighed. The BF:BW ratio was determined and plotted for each bird. Horizontal lines represent the mean and error bars represent the standard error of the mean (SEM).

**Supplementary Figure 3.**
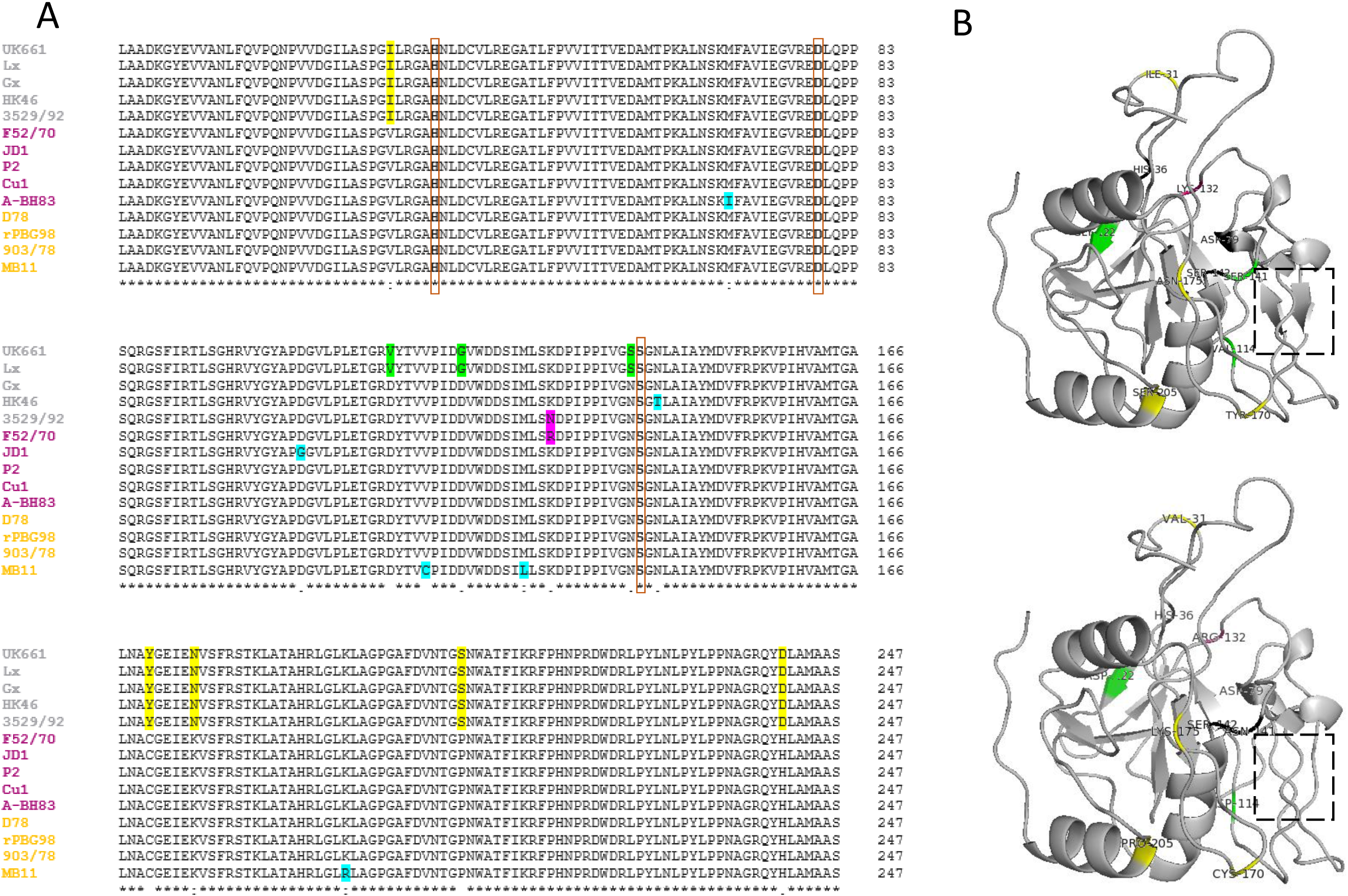
There are nine amino acids different between the UK661 and the F52/70 VP4 proteins that cause differences to the predicted secondary structure of the molecule. The amino acid sequences of five very virulent (grey), six classical (pink) and four cell-adapted (orange) IBDV strains were aligned by Clustal (GenomeNet) (A). Amino acids highlighted in yellow are unique to vvIBDV strains and those in pink are unique to F52/70 compared to all other strains. Residues highlighted in green are found in some but not all vvIBDV strains, and residues highlighted in blue indicate other amino acid differences in the VP4 sequences. The boxed regions are the catalytic triad responsible for the protease function of VP4. The Predicted UK661 (top) and F52/70 (bottom) VP4 structures based on the Yellowtail Ascites Virus VP4 (template 4izk.2.A) modelled using PyMol (B). Mutations between F52/70 and UK661 are labelled in the same colours as indicated in (A) and black sites indicate the location of the catalytic triad responsible for protease function. The dashed-boxed region shows examples of altered secondary structure, with β-strands present in the UK661 VP4 molecule that are not seen in the F52/70 VP4 molecule.

